# The hemagglutinin proteins of clades 1 and 2.3.4.4b H5N1 highly pathogenic avian influenza viruses exhibit comparable attachment patterns to avian and mammalian tissues

**DOI:** 10.1101/2025.06.02.657404

**Authors:** Bingkuan Zhu, Kevin Fung, Hailey Huiyi Feng, Julia A Beatty, Fraser Hill, Anne CN Tse, Christopher J Brackman, Thomas HC Sit, Agnès Poujade, Nicolas Gaide, Mariette Ducatez, Gilles Foucras, Malik Peiris, Shih-Chieh Ti, John M Nicholls, Hui-Ling Yen

## Abstract

The global spread of the A/goose/Guangdong/1/96-lineage H5N1 highly pathogenic avian influenza (HPAI) viruses is accompanied by an expanded host range and the establishment of sustained viral transmission among dairy cattle. To evaluate if the evolving H5N1 viruses have changed tissue tropism over time, we compared the binding patterns of recombinant hemagglutinin (HA) proteins derived from clade 1 (A/Vietnam/1203/04, H5VN) and circulating clade 2.3.4.4b viruses detected from a wild bird (A/Eurasian Teal/Hong Kong/AFCD-HKU-23– 14009–01020/2023, H5HK) and dairy cattle (A/bovine/Ohio/B24OSU-439/2024, H5OH). The HA protein of A(H1N1)pdm09 virus was included for comparison. Using bio-layer interferometry, H1 protein preferentially bound to the α2,6-linked sialoside 6’SLNLN while H5 proteins preferentially bound to the α2,3-linked sialoside 3’SLN. H5OH showed higher binding affinity to 3’SLN than H5HK and H5VN. The attachment pattern of H1 and H5 proteins to the respiratory tissues of different species and dairy cattle mammary glands were evaluated. Compared to the H1 protein, H5 proteins showed stronger binding to the lung epithelial cells of cat, cattle, chicken, ferret, human, and pig, and the clade 2.3.4.4b H5 proteins exhibited increased binding to pig and cattle bronchial epithelial cells. All H5 proteins attached to the alveolar and cistern epithelial cells in mammary glands where α2,3-linked and α2,6-linked sialyl glycans were detected by *Maackia amurensis* lectin II and *Sambucus Nigra* Lectin, respectively. Taken together, the HA proteins of clade 1 and 2.3.4.4b H5N1 viruses generally share comparable attachment patterns to avian and mammalian tissues, despite of evolving into antigenically distinct clades over the past 3 decades.

**IMPORTANCE:** The outbreaks of H5N1 HPAI among US dairy cattle since 2024 have raised concerns of the potential changes in HA receptor binding specificity and tissue tropism. Using insect-cell expressed recombinant HA proteins derived from clade 1 and circulating clade 2.3.4.4b H5N1 viruses, we showed that the dairy cattle H5 protein retained binding specificity for the avian-like α2,3-linked sialoside 3’SLN over the human-like α2,6-linked sialoside 6’SLNLN, with higher binding affinity to 3’SLN than the other H5 proteins. Clade 1 and 2.3.4.4b H5 proteins showed comparable attachment patterns to the mammary tissues of lactating dairy cattle, which showed high expression of α2,3-linked and α2,6-linked sialyl glycans. All H5 proteins also showed comparable attachment patterns to the lungs of cat, cattle, chicken, ferret, human, and pig. Our results suggest that the recent H5N1 outbreaks in dairy cattle may be related to ecological factors rather than changes in HA receptor binding specificity.

## INTRODUCTION

Since the emergence of the A/goose/Guangdong/1/96 (Gs/Gd)-lineage of H5N1 highly pathogenic avian influenza virus (HPAI) from Southern China three decades ago, the hemagglutinin (HA) protein has evolved into multiple antigenically distinct clades, and the Gs/Gd-like viruses have spread across continents via migratory birds (1). The emergence and the expanded geographic distribution of clade 2.3.4.4b H5N1 viruses since 2020 has been accompanied with increasing numbers of outbreaks in domestic and wild bird species (2), spillover infections in humans and a wide range of mammalian species (3, 4), and the establishment of sustained viral transmission among dairy cattle, a new mammalian host for influenza A viruses (5–7).

Since March 2024, multiple States in US have reported outbreaks of genotype B3.13 clade 2.3.4.4b H5N1 virus in dairy cattle (8). Infected cattle presented loss of appetite, massive drop in milk production, and mild respiratory signs (9). Field studies detected higher viral loads in the milk and mammary gland than those detected in the nasal swabs or lungs of infected dairy cattle (6). Experimental studies that used intra-nasal or intra-mammary gland inoculation routes also demonstrated preferential viral replication in the mammary glands over the respiratory tissues (9, 10). Importantly, dairy cattle with intra-mammalian inoculation of a genetically distinct clade 2.3.4.4b virus isolated from a wild goose in Europe (genotype euDG) showed comparable clinical sign as those inoculated with a genotype B3.13 dairy cattle isolate, suggesting that the ability to infect dairy cattle may be shared among clade 2.3.4.4b H5N1 viruses (9), a finding supported by the detection of genotype D1.1 of clade 2.3.4.4b in dairy cattle in January 2025 (11). These findings suggest that H5N1 outbreaks in dairy cattle may be attributed to both virological factor (eg. ability to replicate in mammary gland tissue of dairy cattle) and ecological factors (eg. shared milking devices and inter-state cattle movements). However, it is not clear if early Gs/Gd-lineage H5N1 viruses also possess the capacity to infect dairy cattle. The receptor binding profile of HA proteins determines the host range and cell tropism of influenza A viruses. The HA proteins of avian influenza viruses preferentially bind to α2,3-linked sialyl glycans while the HA proteins of human and swine influenza viruses preferentially bind to α2,6-linked sialyl glycans (2, 3). Here, using insect cell-expressed recombinant HA proteins, we compared the HA attachment pattern of clades 1 and 2.3.4.4b viruses to the mammary tissue of dairy cattle as well as the respiratory tissues of cat, cattle, chicken, ferret, human, and pig.

## RESULTS

### Binding affinity of insect cell-expressed recombinant HA proteins to 3’SLN and 6’SLNLN by bio-infermetry assay

To compare the attachment pattern of HA protein of past and circulating H5N1 viruses, we expressed the HA proteins of A/Vietnam/1203/2004 (clade 1, abbreviated as H5VN), A/Eurasian Teal/Hong Kong/AFCD-HKU-23–14009–01020/2023 (clade 2.3.4.4b, abbreviated as H5HK), and A/bovine/Ohio/B24OSU-439/2024 (clade 2.3.4.4b, abbreviated as H5OH). For comparison, the HA protein from A/California/04/2009 A(H1N1)pmd09 virus was also expressed. The expressed HA proteins were evaluated for their binding affinity to 3’SLN and 6’SLNLN using BLI. All H5 proteins exhibited preferential binding to 3’SLN over 6’SLNLN, while the H1CA exhibited preferential binding to 6’SLNLN over 3’SLN (Fig. 1A). Further analysis using serially 2-fold diluted HA proteins (from 200 to 6.25 nM) to determine binding affinity, we observed that H5OH exhibited higher binding affinity (mean K_D_ ± SD = 18.55 ± 1.35 nM) than H5VN (36.45 ± 1.75 nM) and H5HK (46.95 ± 3.45 nM) (One-way ANOVA, p<0.01) (Fig. 1B). While the clade 1 H5VN differed by the clade 2.3.4.4b H5OH by 40 amino acids, the two clade 2.3.4.4.b H5HK and H5OH only differed by 4 amino acids (L111M, L122Q, T199I, V214A, H3 numbering) in HA1 (Fig. 1C). Specifically, the T199I change have been reported to increase the HA biding breath to α2,3-linked *N-*acetyllactosamines of the first human H5N1 isolate (A/Texas/37/2024) reported in the dairy cattle outbreak (12).

**Figure 1.**
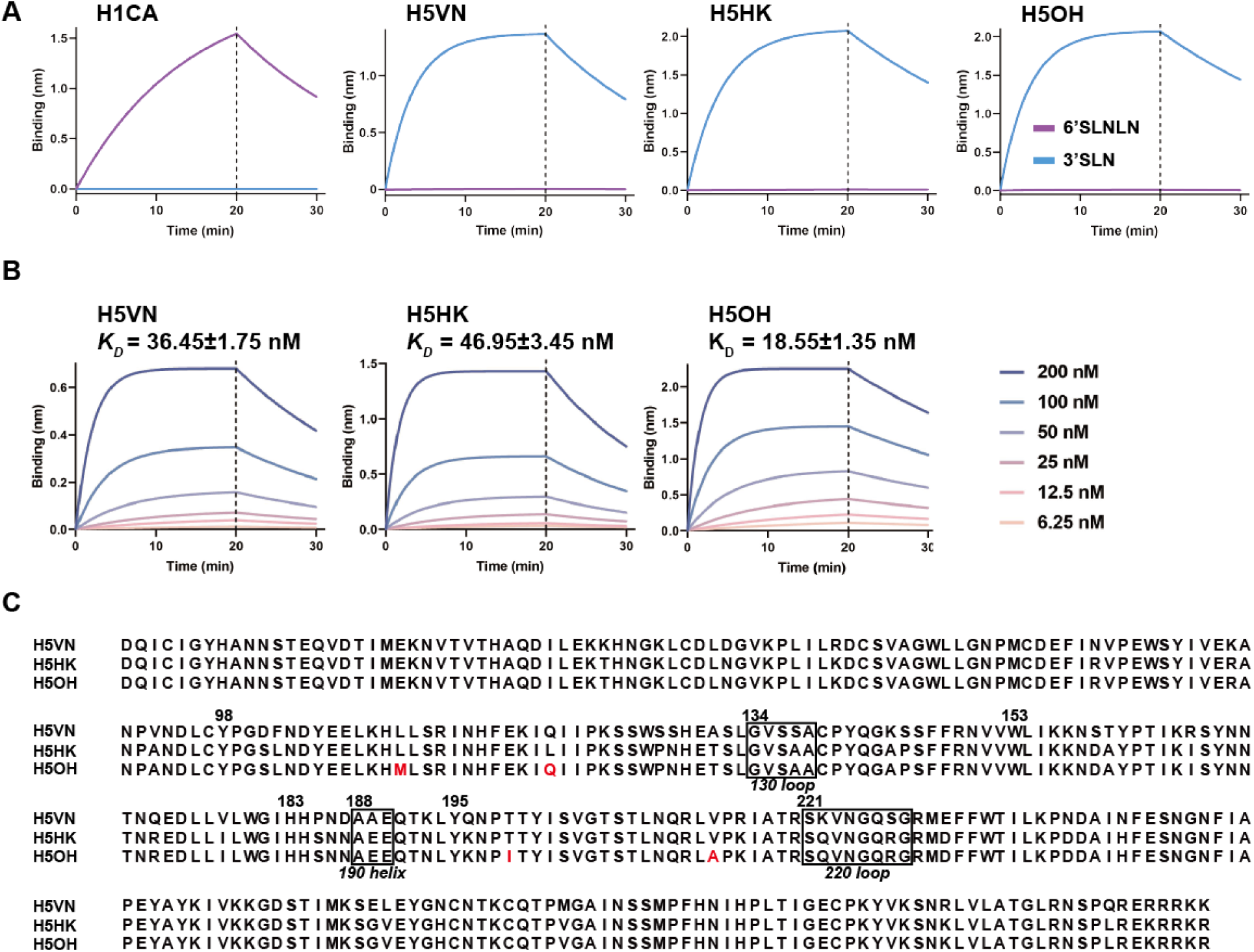
Binding kinetics of recombinant HA proteins to α2,3-linked and α2,6-linked sialyl glycans using BLI. (A) Streptavidin biosensors were immobilized with biotinylated α2,3-linked (3’SLN) or α2,6-linked (6’SLNLN) glycans at the concentration of 1 μg/mL, followed by incubation with recombinant HA proteins 67.5 nM for 20 minutes at 30°C. Binding signal over time are shown. (B) Kinetics of recombinant HA binding with 3’SLN measured by BLI. Curves were generated with equal densities of immobilized 3’SLN and the indicated purified HA protein concentrations. (C) Amino acid alignment of HA1 from H5 viruses in this study. Residues that differ between H5HK and H5OH were marked in red.

### Clade 1 and clade 2.3.4.4b H5 proteins bind to the mammary tissues of lactating cows

Infection of mammary tissues has been a unique feature observed from the outbreaks of clade 2.3.4.4b H5N1 in dairy cattle (6, 9, 10). We compared the attachment pattern of H5VN, H5HK, H5OH, and H1CA to the mammary gland tissues of lactating cows (Fig. 2A). With recombinant proteins diluted to 12.5 μg/mL, we observed no binding of H1CA, but all three H5 recombinant proteins showed comparable binding intensity to the alveolar and cistern epithelial cells (Fig. 2B). It is noteworthy that no binding to the ductal epithelial cells was observed (Fig. 2B). To further characterize α2,3 and α2,6-linked sialyl glycans presented in the mammary tissues, we performed lectin staining with SNA that preferentially binds to NeuAcα2,6Galβ1,4GlcNAc, MAL-I that preferentially binds to NeuAcα2,3Galβ1,4GlcNAc in N-glycans or O-glycans, and MAL-II that preferentially binds to NeuAcα2,3Galβ1,3GalNAc in O-glycans. Both SNA and MAL-II showed apparent binding to the alveolar epithelial cells and minor binding to the cistern epithelial cells. Interestingly, no apparent MAL-I binding was observed, suggesting that the mammary tissue of lactating cows may express low level of NeuAcα2,3Galβ1,4GlcNAc (Fig. 2A).

**Fig. 2.**
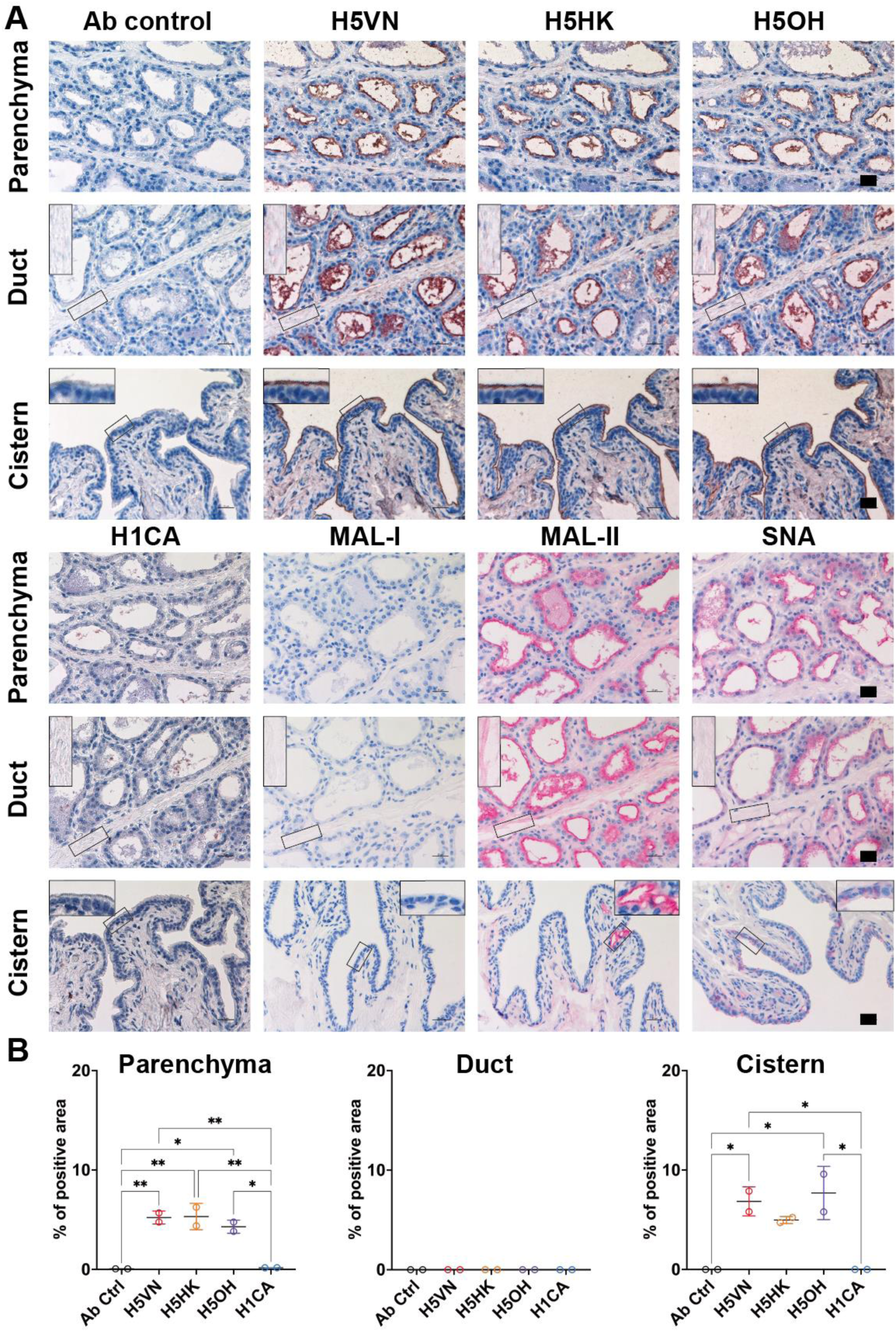
Attachment pattern of HA proteins and lectins to bovine mammary tissues. (A) Recombinant HA proteins (12.5 μg/mL) and plant lectins (20 μg/mL for SNA/MAL-I, and 10 μg/mL for MAL-II) were evaluated for their attachment pattern to formalin-fixed mammary gland tissues of lactating dairy cattle using protein histochemistry. The nuclei were counterstained with hematoxylin (blue). Inserts are digital magnifications of the boxed area. Scale bar indicates 25 μm. (B) The binding intensity of HA proteins were quantified using Qupath (version 0.5.1) from two independently repeated slides. The HA binding intensity to ductal and cistern epithelial cells were measured using the sub-panels showing the specific signals from the epithelial cells. One-way ANOVA was used to compare binding intensity of different HA proteins. *, p<0.05; **, p<0.01.

### H5 proteins exhibited strong binding to lung epithelial cells of chicken, cat, cattle, ferret, human, and pig

We further evaluated the attachment pattern of recombinant HA proteins to the respiratory tissues of different species. All H5 protein binds to chicken trachea and lung epithelial cells while no binding of H1CA was observed (Fig. 3A and 3B). The recombinant HA proteins also exhibited different attachment pattern to the mammalian respiratory tissues (Fig. 4 and 5). In the ferret bronchus, H1CA showed patchy but stronger binding to the bronchial epithelial cells than the H5 proteins, while the clade 2.3.4.4b H5 proteins showed stronger binding to pig and cattle bronchus than H1CA (Fig. 4A and 4B). All three H5 recombinant proteins showed stronger binding to the lung epithelial cells of cat, cattle, human, and pig than the H1 protein, with the H5OH exhibiting stronger binding to the ferret lung epithelial cells than H5HK or H5VN (Fig. 5A and 5B). Taken together, the results suggest that the HA proteins derived from clade 1 and clade 2.3.4.4b H5N1 viruses showed minor differences in their attachment patterns to the respiratory tissues of different species.

**Fig 3.**
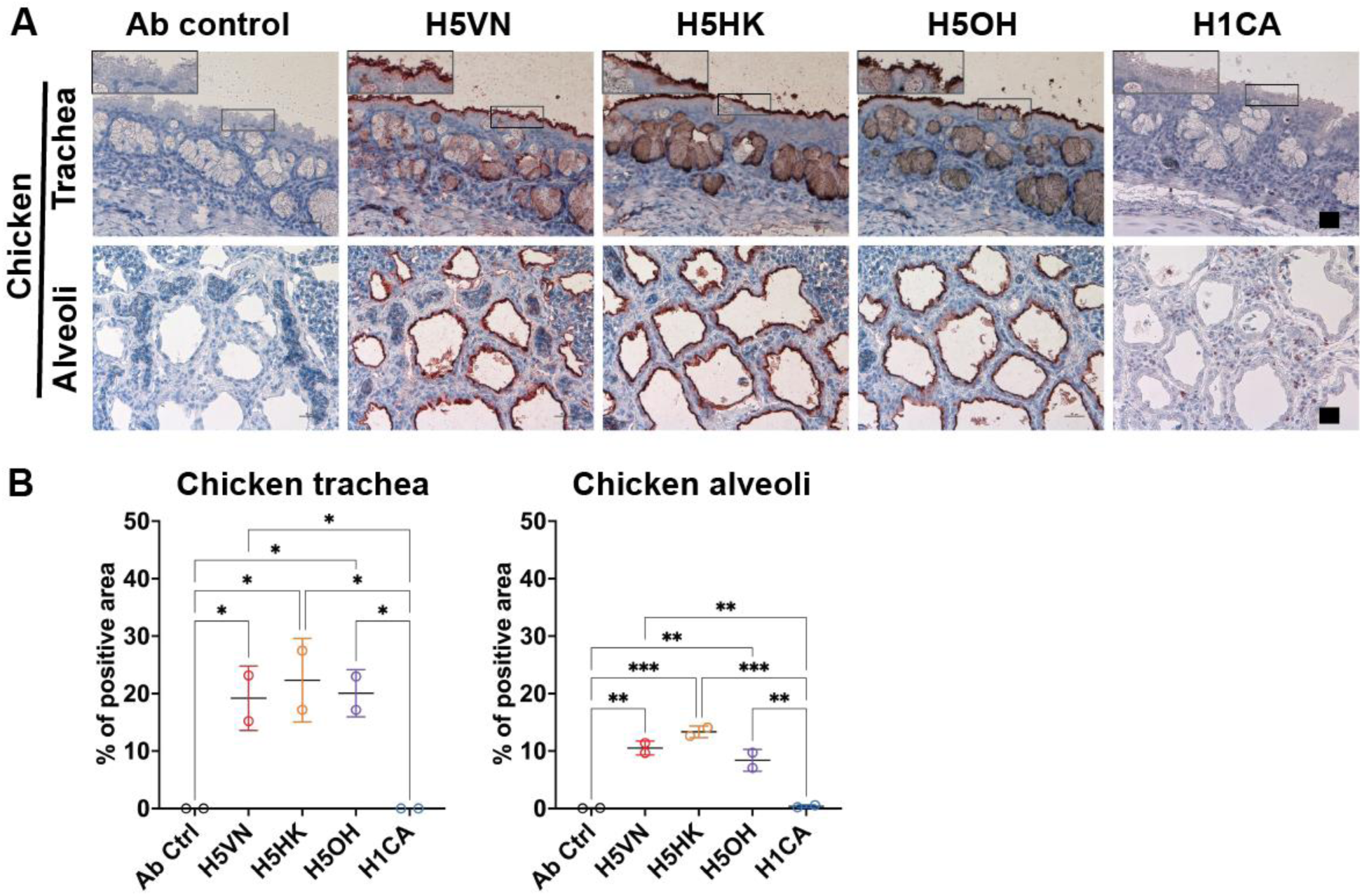
Attachment pattern of HA proteins to chicken respiratory tissues. (A) Recombinant HA proteins were diluted to 12.5 μg/mL and incubated with to formalin-fixed chicken trachea and lungs using protein histochemistry. The nuclei were counterstained with hematoxylin (blue). Inserts are digital magnifications of the boxed area. Scale bar indicates 25 μm. (B) The binding intensity of HA proteins were quantified using Qupath (version 0.5.1) from two independently repeated slides. One-way ANOVA was used to compare binding intensity of different HA proteins. *, p<0.05; **, p<0.01; ***, p<0.001.

**Fig 4.**
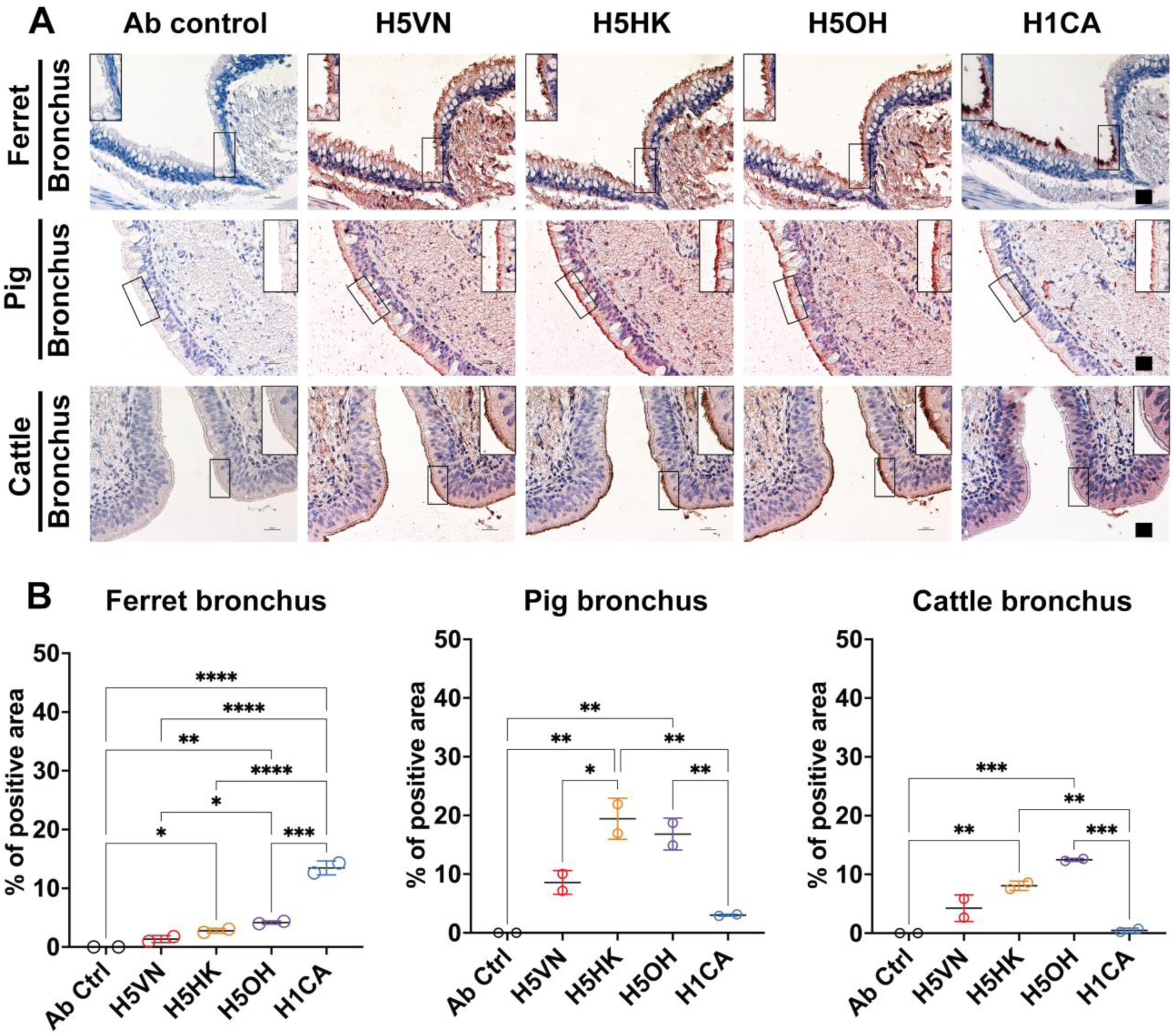
Attachment pattern of HA proteins to mammalian bronchial epithelial cells. (A) Recombinant HA proteins were diluted to 12.5 μg/mL and incubated to formalin-fixed bronchus from cattle, ferret and pig using protein histochemistry. The nuclei were counterstained with hematoxylin (blue). Inserts are digital magnifications of the boxed area. Scale bar indicates 25 μm. (B) The binding intensity of HA proteins were quantified using Qupath (version 0.5.1) from two independently repeated slides, with measurements of the sub-panels showing the bronchial epithelial cells. One-way ANOVA was used to compare binding intensity of different HA proteins. *, p<0.05; **, p<0.01; ***, p<0.001; ****<0.0001.

**Fig 5.**
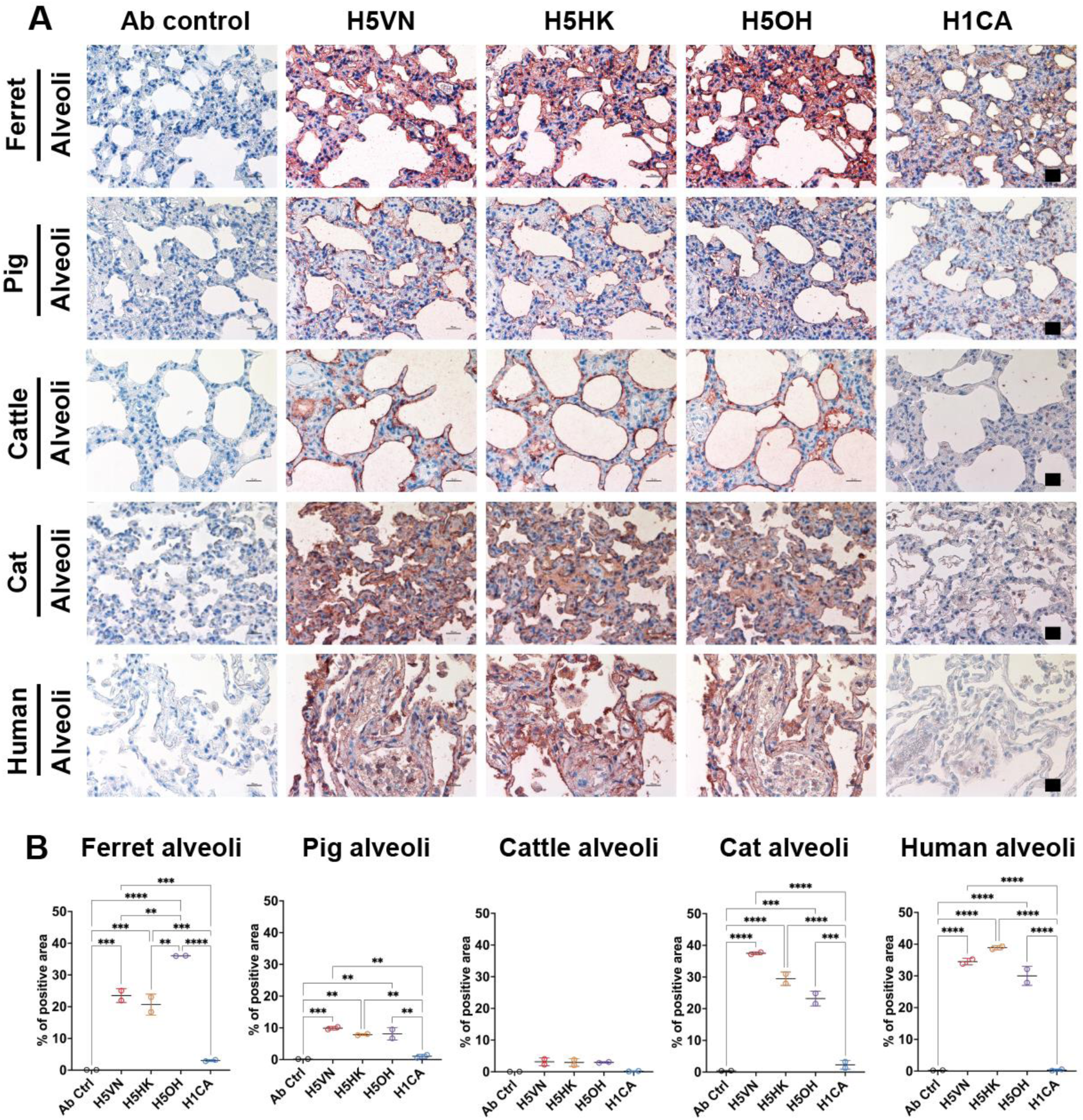
Attachment pattern of HA proteins to mammalian lung alveolar epithelial cells. (A) Recombinant HA proteins were diluted to 12.5 μg/mL and incubated to formalin-fixed lungs tissues from cat, cattle, ferret, human, and pig using protein histochemistry. The nuclei were counterstained with hematoxylin (blue). (B) The binding intensity of HA proteins were quantified using Qupath (version 0.5.1) from two independently repeated slides. One-way ANOVA was used to compare binding intensity of different HA proteins. **, p<0.01; ***, p<0.001; ****<0.0001.

## DISCUSSION

The expanded host range and the sustained transmission of clade 2.3.4.4b H5N1 viruses among dairy cattle have raised concerns on the potential changes in the HA attachment pattern to sialyl glycans expressed on animal tissues, which may affect viral transmissibility and pandemic potential. Our results suggest that H5 proteins derived from clade 1 and clade 2.3.4.4b generally share comparable attachment pattern to the cattle mammary tissues and the respiratory tissues of different hosts, suggesting that the H5N1 outbreaks in dairy cattle may be related to ecological factors rather than changes in HA receptor binding specificity. Additional epidemiological studies and environmental sampling are needed to identify risk factors associated with the introduction of H5N1 into dairy herds.

Using BLI, we observed that all H5 proteins preferentially bind to 3’SLN with no detectable binding to 6’SLNLN. H5OH also showed higher binding affinity to 3’SLN than H5HK or HKVN. The BLI assay is sensitive and quantitative but only allows evaluating HA binding to specific glycans each time. On the other hand, the use of recombinant HA proteins allowed assessing viral tropism in tissues presented with diverse glycan structures, although this method is more qualitative than quantitative. Using recombinant H5 protein derived from clade 1, clade 2.1 and an early clade 2.3.4.4b strain isolated in 2016, Ríos Carrasco reported H5 proteins may bind to the lung alveolar epithelial cells but not to the tracheal epithelial cells of cattle, pig, and horse (13). Here, we showed that H5 proteins derived from clade 1 and circulating clade 2.3.4.4b viruses may bind to the bronchial epithelial cells of ferret, pig, and cattle, albeit with different binding intensity. Specifically, H1 protein showed better binding than H5 proteins to the ferret bronchial epithelial cells. However, the two clade 2.3.4.4b H5 proteins showed better binding to the pig and cattle bronchial epithelial cells than the H1 protein. This suggests that pigs may be susceptible to infection of clade 2.3.4.4b viruses (14) and co-infection with other swine influenza viruses may pose a risk for the emergence of novel reassortant viruses. Bauer et al. also reported that a clade 2.3.4.4b virus (A/Capsian gull/Netherlands/1/2022) showed increased replication and attachment to human tracheal/bronchial epithelial cells than a clade 2.1 virus A/Indonesia/5/2005 (15). In addition, we observed that the clade 1 and 2.3.4.4b H5 proteins showed comparable attachment to the lung alveolar epithelial cells of chicken, cat, cattle, human, and pig; however, the H5OH showed increased binding to ferret lung alveolar epithelial cells than H5VN or H5HK. Taken together, our results are consistent with previous studies that the HA protein of cattle H5N1 viruses retained predominantly avian-type receptor binding specificity (7, 12, 13, 16–19), although few studies reported that the cattle H5N1 virus may bind to human-type receptors (7, 20, 21). Variations in the quantity of HA proteins and HA valency used by different assays may have affected the results between different studies.

Experimental infection showed that the mammary gland may serves as the main site of replication in dairy cattle by the clade 2.3.4.4b H5N1 viruses (9, 10). Since clade 1 and clade 2.3.4.4b H5 proteins all bind to the alveolar and cistern epithelial cells in the mammary glands, it is likely that clade 1 H5N1 virus may similarly cause infection in mammary gland tissue given the proper opportunity. The attachment pattern of H5 proteins in mammary glands are in accordance with the results reported by previous studies (7, 13). Using lectin staining, we observed predominant expression of NeuAcα2,6Galβ1,4GlcNAc (detected by SNA) and NeuAcα2,3Galβ1,3GalNAc (detected by MAL-II) in the mammary gland tissues of lactating dairy cattle, while the expression of NeuAcα2,3Galβ1,4GlcNAc (detected by MAL-I) was low. This result in combination of the H5 protein attachment pattern suggest H5 proteins bind to the NeuAcα2,3Galβ1,3GalNAc O-glycans in the mammary tissues (12). Interestingly, although α2,6-linked sialyl glycans were distributed along the gland alveolar and cistern epithelial cells, we did not observe any binding of the H1 protein to these tissues. Our result is consistent with those reported previously, including limited attachment of recombinant H1 protein derived from a mouse-adapted influenza strain A/Puerto Rico/8/34 to dairy cattle mammary tissues (13), and restricted replication of an A(H1N1)pdm09 virus in the *ex vivo* culture of bovine mammary gland (22).

Taken together, the data available to date support that the clade 2.3.4.4b retained comparable receptor binding profile as the early H5N1 viruses. However, it’s important to note that H5N1 viruses continue to cause spillover infections in mammals, which provide opportunities for viral adaptation. A study reported that a single Gln226Leu mutation may switch the cattle H5N1 virus binding specificity to human-type receptors (19). Continuous efforts on surveillance and monitoring of the evolution of H5N1 viruses isolated from different host species is essential for pandemic preparedness.

## MATEIRALS AND METHODS

### Expression of recombinant HA proteins

Soluble recombinant HA proteins were expressed and purified following established protocols (23, 24). Genes encoding the ectodomains of the HA protein of the A(H1N1)pdm09 and A(H5N1) (with the multibasic cleavage site removed) viruses were subcloned into the baculovirus transfer vector pFastBac1 (Invitrogen), in frame with an N-terminal gp67 signal peptide, a C-terminal trimerization foldon sequence from bacteriophage T4, followed by a thrombin cleavage site and a His_6_-tag. Transfection and baculovirus amplification were performed in Sf9 cells using the Bac-to-Bac baculovirus expression system (Invitrogen). On day 3 post-infection, supernatants were harvested, and the HA proteins were purified using Ni-charged immobilized metal affinity chromatography (IMAC) resin (Bio-Rad). Recombinant proteins were stored in PBS with 20% sucrose (Sigma) in aliquots at -80°C.

### HA binding affinity to glycans using bio-layer interferometry (BLI)

The expressed recombinant HA proteins were evaluated for their binding affinity to biotinylated 3’SLN (Neu5Acα2-3Galβ1-4GlcNAcβ-sp3-PAA-biot) (GlycoNZ) and biotinylated 6’SLNLN (Neu5Acα2-6Galβ1-4GlcNAcβ1-3Galβ1-4GlcNAcβ-sp3-PAA-biot) (GlycoNZ) using BLI. Assays were performed using the Octet Red96e system (FortéBio) in 96-well microplates as previously described (25). Dulbecco’s PBS with calcium, magnesium, and 0.005% Tween-20 was used as the assay buffer to reconstitute protein and glycan molecules. Biotinylated 3’SLN and 6’SLNLN (GlycoNZ) were preloaded to the streptavidin-coated biosensors (FortéBio) at 1 μg/mL for 10 min. HA proteins diluted to 67.5 nM were pre-conjugated with the mouse anti-His-tag antibody (Thermo Scientific, clone # MA1-21315) and the horseradish peroxidase (HRP)-conjugated goat anti-mouse secondary antibody (Abcam, clone # ab6789) at the molar ratio of 2:1:2 for 30 min on ice before added to the wells. The 96-well plates were incubated at 30 °C for 30 minutes, with sample plates agitated at 1000 RPM. Binding kinetics of H5 proteins to 3’SLN were measured for 20 minutes at HA concentrations of 6.25 nM, 12.5 nM, 25 nM, 50 nM, 100 nM, and 200 nM. The data were fitted with a 1:1 binding model to evaluate HA binding affinity (K_D_) to 3’SLN.

### Immunohistochemical staining with recombinant HA proteins

Formalin-fixed and paraffin-embedded tissue blocks and sections were prepared by the Department of Pathology, Li Ka Shing Faculty of Medicine, The University of Hong Kong or obtained from collaborating laboratories. Immunohistochemistry was performed according to a previously described protocol (13) with minor modifications. Briefly, the sections were dried at 60°C for 20 min, deparaffinized, and rehydrated. Antigen retrieval was achieved by boiling the sections in 10 mM sodium citrate (pH 6.0) for 20 min. Endogenous peroxidase activity was quenched using 3% hydrogen peroxidase, and nonspecific binding was blocked with 10% goat serum (Giobco). Histidine-tagged recombinant HA proteins were diluted to 12.5 μg/mL and were pre-conjugated with the mouse anti-His-tag primary antibody and the HRP-conjugated goat anti-mouse secondary antibody (Abcam) as described. This pre-complexed mixture was then applied to the tissue sections and incubated for 2 hours at room temperature followed by washing with PBS buffer containing 0.05% (v/v) Tween-20 (PBST). The chromogen 3-amino-9-ethylcarbazole (AEC) (Sigma-Aldrich) was used as the HRP substrate. Tissues were counterstained with Gill’s hematoxylin (Vector Laboratories), mounted with permanent aqueous mounting medium (Bio-Rad) and examined using a Nikon Eclipse Ti-S microscope. The experiments were repeated independently twice.

### Lectin staining

To evaluate the distribution of α2,3-linked and α2,6-linked sialyl glycans in tissue slides, biotinylated *Sambucus Nigra* Lectin (SNA), *Maackia amurensis* lectin I (MAL-I), *Maackia amurensis* lectin II (MAL-II) were used for staining. SNA is known to preferentially bind to α2,6-linked terminal sialic acids (SA) (Neu5Acα2,6Galβ1,4GlcNAc). MAL-I and MAL-II preferentially binds to α2,3-linked terminal SA, but MAL-I preferred Neu5Acα2,3Galβ1,4GlcNAc while MAL-II preferred Neu5Acα2,3Galβ1,3GalNAc. Antigen retrieval and endogenous peroxidase blocking were performed as described above. After blocking with 0.1% Bovine Serum Albumin (Sigma-Aldrich), the slides were incubated with 20 μg/mL SNA/MAL-I, or 10 μg/mL MAL-II at room temperature for 1 hour, followed by incubation with alkaline phosphatase-conjugated streptavidin (Vector Laboratories) for 45 minutes. The slides were then developed with Vector Red Substrate Kit (Vector Laboratories). After counterstaining with Gill’s hematoxylin (Vector Laboratories), the slides were counterstained with Scott’s tap water (Sigma-Aldrich), air-dried and mounted with Permount (Fisher Scientific).

### Statistical analysis

To compare HA binding intensity to the respiratory and mammary tissues, the chromagen AEC signal was quantified using the Qupath software (26) from two independently stained tissue slides as shown. One-way ANOVA test was used to compare binding signal of different recombinant HA proteins.

## ACKNOWLEDGEMENTS

This study was supported by NIAID, National Institutes of Health, USA (Contract 75N93021C00016) and RGC Theme-based Research Schemes, Hong Kong SAR, China (T11-712/19-N).

## REFERENCES

1. Bodewes R, Kuiken T. 2018. Changing Role of Wild Birds in the Epidemiology of Avian Influenza A Viruses. Adv Virus Res 100:279–307.

2. Adlhoch C, Fusaro A, Gonzales JL, Kuiken T, Marangon S, Mirinaviciute G, Niqueux É, Stahl K, Staubach C, Terregino C, Broglia A, Baldinelli F. 2023. Avian influenza overview December 2022 - March 2023. Efsa j 21:e07917.

3. Plaza PI, Gamarra-Toledo V, Euguí JR, Lambertucci SA. 2024. Recent Changes in Patterns of Mammal Infection with Highly Pathogenic Avian Influenza A(H5N1) Virus Worldwide. Emerg Infect Dis 30:444–452.

4. WHO. 20 January 2025 2025. Cumulative number of confirmed human cases for avian influenza A(H5N1) reported to WHO, 2003-2025, 20 January 2025. https://www.who.int/publications/m/item/cumulative-number-of-confirmed-human-cases-for-avian-influenza-a(h5n1)-reported-to-who--2003-2025--20-january-2025. Accessed on 30 May 2025.

5. Burrough ER, Magstadt DR, Petersen B, Timmermans SJ, Gauger PC, Zhang J, Siepker C, Mainenti M, Li G, Thompson AC, Gorden PJ, Plummer PJ, Main R. 2024. Highly Pathogenic Avian Influenza A(H5N1) Clade 2.3.4.4b Virus Infection in Domestic Dairy Cattle and Cats, United States, 2024. Emerg Infect Dis 30:1335–1343.

6. Caserta LC, Frye EA, Butt SL, Laverack M, Nooruzzaman M, Covaleda LM, Thompson AC, Koscielny MP, Cronk B, Johnson A, Kleinhenz K, Edwards EE, Gomez G, Hitchener G, Martins M, Kapczynski DR, Suarez DL, Alexander Morris ER, Hensley T, Beeby JS, Lejeune M, Swinford AK, Elvinger F, Dimitrov KM, Diel DG. 2024. Spillover of highly pathogenic avian influenza H5N1 virus to dairy cattle. Nature 634:669–676.

7. Song H, Hao T, Han P, Wang H, Zhang X, Li X, Wang Y, Chen J, Li Y, Jin X, Duan X, Zhang W, Bi Y, Jin R, Sun L, Wang N, Gao GF. 2025. Receptor binding, structure, and tissue tropism of cattle-infecting H5N1 avian influenza virus hemagglutinin. Cell 188:919–929.e9.

8. CDC. 2025. Current Situation: Bird Flu in Dairy Cows. https://www.cdc.gov/bird-flu/situation-summary/mammals.html. Accessed 30 May 2025.

9. Halwe NJ, Cool K, Breithaupt A, Schön J, Trujillo JD, Nooruzzaman M, Kwon T, Ahrens AK, Britzke T, McDowell CD, Piesche R, Singh G, Pinho Dos Reis V, Kafle S, Pohlmann A, Gaudreault NN, Corleis B, Ferreyra FM, Carossino M, Balasuriya UBR, Hensley L, Morozov I, Covaleda LM, Diel DG, Ulrich L, Hoffmann D, Beer M, Richt JA. 2025. H5N1 clade 2.3.4.4b dynamics in experimentally infected calves and cows. Nature 637:903–912.

10. Baker AL, Arruda B, Palmer MV, Boggiatto P, Sarlo Davila K, Buckley A, Ciacci Zanella G, Snyder CA, Anderson TK, Hutter CR, Nguyen TQ, Markin A, Lantz K, Posey EA, Kim Torchetti M, Robbe-Austerman S, Magstadt DR, Gorden PJ. 2025. Dairy cows inoculated with highly pathogenic avian influenza virus H5N1. Nature 637:913–920.

11. USDA. 2025. APHIS Confirms D1.1 Genotype in Dairy Cattle in Nevada. https://www.aphis.usda.gov/news/program-update/aphis-confirms-d11-genotype-dairy-cattle-nevada-0. Accessed 30 May 2025.

12. Good MR, Fernandez-Quintero ML, Ji W, Rodriguez AJ, Han J, Ward AB, Guthmiller JJ. 2024. A single mutation in dairy cow-associated H5N1 viruses increases receptor binding breadth. Nat Commun 15:10768.

13. Ríos Carrasco M, Gröne A, van den Brand JMA, de Vries RP. 2024. The mammary glands of cows abundantly display receptors for circulating avian H5 viruses. J Virol 98:e0105224.

14. Kwon T, Trujillo JD, Carossino M, Lyoo EL, McDowell CD, Cool K, Matias-Ferreyra FS, Jeevan T, Morozov I, Gaudreault NN, Balasuriya UBR, Webby RJ, Osterrieder N, Richt JA. 2024. Pigs are highly susceptible to but do not transmit mink-derived highly pathogenic avian influenza virus H5N1 clade 2.3.4.4b. Emerg Microbes Infect 13:2353292.

15. Bauer L, Leijten L, Iervolino M, Chopra V, van Dijk L, Power M, Spronken M, Rijnink W, Funk M, de Vries RD, Richard M, Kuiken T, van Riel D. 2024. A 2022 avian H5N1 influenza A virus from clade 2.3.4.4b attaches to and replicates better in human respiratory epithelium than a 2005 H5N1 virus from clade 2.3.2.1. bioRxiv doi:10.1101/2024.11.27.625596:2024.11.27.625596.

16. Chopra P, Ray SD, Page CK, Shepard JD, Kandeil A, Jeevan T, Bowman AS, Ellebedy AH, Webby RJ, de Vries RP, Tompkins SM, Boons GJ. 2025. Receptor-binding specificity of a bovine influenza A virus. Nature 640:E21–e27.

17. Santos JJS, Wang S, McBride R, Adams L, Harvey R, Zhao Y, Wrobel AG, Gamblin S, Skehel J, Lewis NS, Paulson JC, Hensley SE. 2025. Bovine H5N1 binds poorly to human-type sialic acid receptors. Nature 640:E18–e20.

18. Yang J, Qureshi M, Kolli R, Peacock TP, Sadeyen JR, Carter T, Richardson S, Daines R, Barclay WS, Brown IH, Iqbal M. 2025. The haemagglutinin gene of bovine-origin H5N1 influenza viruses currently retains receptor-binding and pH-fusion characteristics of avian host phenotype. Emerg Microbes Infect 14:2451052.

19. Lin TH, Zhu X, Wang S, Zhang D, McBride R, Yu W, Babarinde S, Paulson JC, Wilson IA. 2024. A single mutation in bovine influenza H5N1 hemagglutinin switches specificity to human receptors. Science 386:1128–1134.

20. Eisfeld AJ, Biswas A, Guan L, Gu C, Maemura T, Trifkovic S, Wang T, Babujee L, Dahn R, Halfmann PJ, Barnhardt T, Neumann G, Suzuki Y, Thompson A, Swinford AK, Dimitrov KM, Poulsen K, Kawaoka Y. 2024. Pathogenicity and transmissibility of bovine H5N1 influenza virus. Nature 633:426–432.

21. Gu C, Maemura T, Guan L, Eisfeld AJ, Biswas A, Kiso M, Uraki R, Ito M, Trifkovic S, Wang T, Babujee L, Presler R, Jr., Dahn R, Suzuki Y, Halfmann PJ, Yamayoshi S, Neumann G, Kawaoka Y. 2024. A human isolate of bovine H5N1 is transmissible and lethal in animal models. Nature 636:711–718.

22. Imai M, Ueki H, Ito M, Iwatsuki-Horimoto K, Kiso M, Biswas A, Trifkovic S, Cook N, Halfmann PJ, Neumann G, Eisfeld AJ, Kawaoka Y. 2025. Highly pathogenic avian H5N1 influenza A virus replication in ex vivo cultures of bovine mammary gland and teat tissues. Emerg Microbes Infect 14:2450029.

23. Margine I, Palese P, Krammer F. 2013. Expression of functional recombinant hemagglutinin and neuraminidase proteins from the novel H7N9 influenza virus using the baculovirus expression system. J Vis Exp doi:10.3791/51112:e51112.

24. Shi Y, Zhang W, Wang F, Qi J, Wu Y, Song H, Gao F, Bi Y, Zhang Y, Fan Z, Qin C, Sun H, Liu J, Haywood J, Liu W, Gong W, Wang D, Shu Y, Wang Y, Yan J, Gao GF. 2013. Structures and receptor binding of hemagglutinins from human-infecting H7N9 influenza viruses. Science 342:243–7.

25. Du W, Wolfert MA, Peeters B, van Kuppeveld FJM, Boons GJ, de Vries E, de Haan CAM. 2020. Mutation of the second sialic acid-binding site of influenza A virus neuraminidase drives compensatory mutations in hemagglutinin. PLoS Pathog 16:e1008816.

26. Bankhead P, Loughrey MB, Fernández JA, Dombrowski Y, McArt DG, Dunne PD, McQuaid S, Gray RT, Murray LJ, Coleman HG, James JA, Salto-Tellez M, Hamilton PW. 2017. QuPath: Open source software for digital pathology image analysis. Sci Rep 7:16878.

